# Novel System to Monitor *In Vivo* Neural Graft Activity After Spinal Cord Injury

**DOI:** 10.1101/2021.08.29.458127

**Authors:** Kentaro Ago, Narihito Nagoshi, Kent Imaizumi, Takahiro Kitagawa, Momotaro Kawai, Keita Kajikawa, Reo Shibata, Yasuhiro Kamata, Kota Kojima, Munehisa Shinozaki, Takahiro Kondo, Satoshi Iwano, Atsushi Miyawaki, Masanari Ohtsuka, Haruhiko Bito, Kenta Kobayashi, Shinsuke Shibata, Tomoko Shindo, Jun Kohyama, Morio Matsumoto, Masaya Nakamura, Hideyuki Okano

## Abstract

Expectations for neural stem/progenitor cell (NS/PC) transplantation as a treatment for spinal cord injury (SCI) are increasing. However, whether and how grafted cells are incorporated into the host neural circuit and contribute to motor function recovery remain unknown. The aim of this project was to establish a novel non-invasive *in vivo* imaging system to visualize the activity of neural grafts by which we can simultaneously demonstrate the circuit-level integration between the graft and host, and the contribution of graft neuronal activity to host behaviour. We introduced Akaluc, a newly engineered luciferase, under control of a potent neuronal activity-dependent synthetic promoter, E-SARE, into NS/PCs and engrafted the cells into SCI model mice. Through the use of this system, we reveal that the activity of grafted cells was integrated with host behaviour and driven by host neural circuit inputs. This non-invasive system is expected to help elucidate the therapeutic mechanism of cell transplantation treatment for SCI and determine better therapy techniques that maximize the function of cells in the host circuit.

## Introduction

Spinal cord injury (SCI) results in severe neurological dysfunction, including motor, sensory, and autonomic paralyses. In recent years, many attempts to develop cell transplantation therapies to promote regeneration of the damaged spinal cord have been made. Neural stem/progenitor cells (NS/PCs) are one of the most promising resources for such therapies (Assinck, Duncan, Hilton, Plemel, & Tetzlaff, 2017; Cummings et al., 2005; Iwanami et al., 2005). Several studies have proposed that NS/PC grafts can form neuronal relays across sites of spinal transection (Adler, Lee-Kubli, Kumamaru, Kadoya, & Tuszynski, 2017; Kadoya et al., 2016; Kumamaru et al., 2018; Lu et al., 2012), that is, combine input from the rostral part of the host to the graft and output from the graft to the caudal part, which are thought to play a major role in functional recovery. However, a detailed characterization of neuronal relay has not been carried out, and how the graft functionally integrates into the host neural circuit is poorly understood. This is mainly because no current technologies can directly monitor the relationship between graft cell activity and the behaviours and circuit-level activities of the host. To elucidate functional host-graft coordination and to evaluate how the graft influences host neural circuit activity and host behaviour, a novel non-invasive *in vivo* imaging technique to monitor the activity of graft neurons over time within the living host is needed.

To realize such an *in vivo* monitoring system, we focused on two novel technologies. The first is the AkaBLI system (the combination of the AkaLuc enzyme and AkaLumine-HCl as a substrate with high permeability) (Iwano et al., 2018; Kuchimaru et al., 2016). Bioluminescence imaging (BLI) is a non-invasive method for measuring light output from cells expressing the enzyme luciferase after luciferin (substrate) administration in living animals (Hara-Miyauchi et al., 2012). AkaBLI is a newly developed red-shifted BLI system that produces bright emission spectra and enables deep tissue imaging in living animals (Iwano et al., 2018), which is the most appropriate for widefield non-invasive monitoring of gene expression from graft cells in injured spinal cords. The second is enhanced synaptic activity-responsive element (E-SARE), a potent neuronal activity-dependent synthetic promoter (Kawashima et al., 2013). When a neuron becomes active, it switches on immediate-early genes (IEGs), such as *Fos*, *Arc*, and *Egr1*, even in spinal cord neurons, and the promoters/enhancers of IEGs are used as activity-dependent reporter systems (Bonner et al., 2011; Guzowski, McNaughton, Barnes, & Worley, 1999). Among these promoters is the synthetic promoter E-SARE, which is based on the SARE enhancer element of the *Arc* promoter, and drives a significantly superior neuronal activity-dependent gene expression to any other existing IEG promoters.

In this study, we combined AkaBLI and E-SARE technology and established a novel non-invasive system to visualize the neuronal activity of the graft *in vivo*. We succeeded in imaging the active ensemble dynamics of NS/PC-derived cells grafted in injured spinal cords. Using this system, we confirmed that graft activity is linked to host behaviour and that the host circuit regulates graft activity.

## Results

### The ESAL system: neuron activity monitoring by bioluminescence

To establish a bioluminescence-based system to visualize neuronal activity, we first constructed a lentiviral vector for the expression of AkaLuc, a luciferase optimized for red-shifted bioluminescence, under the control of E-SARE, a potent neuronal activity-dependent promoter generated from *Arc* enhancer elements (Figure 1a). We termed this system ESAL (<u>E</u>-<u>S</u>ARE-<u>A</u>ka<u>L</u>uc). In the ESAL system, we also fused AkaLuc with the Venus protein for simultaneous fluorescent labelling and with the PEST sequence to shorten the half-life of the fusion protein (Li et al., 1998). The Venus protein is an *Aequorea victoria*-derived YFP containing mutations that cause rapid maturation and increased environmental resistance (Nagai et al., 2002). We then transfected the ESAL lentiviral vector into human induced pluripotent stem cell (iPSC)- derived NS/PCs, identified as ESAL-NS/PCs, and induced their differentiation into neurons (Figure 1b–h). When stimulated by a depolarizing concentration of potassium chloride (50 mM), ESAL-NS/PC-derived neurons showed a significant increase in AkaLuc photon count compared with that of un-stimulated Controls (Figure 1b, c). We also detected increases in Venus fluorescence and IEG expression upon 50 mM KCl stimulation (Figure 1d, e). Thus, we confirmed that the ESAL system was highly sensitive to depolarizing stimulation in neurons, while showing little responses in non-neuronal cells (Figure 1f–h). These data suggest that the ESAL system can successfully monitor the neuronal activity of NS/PC-derived neurons.

**Figure 1.**
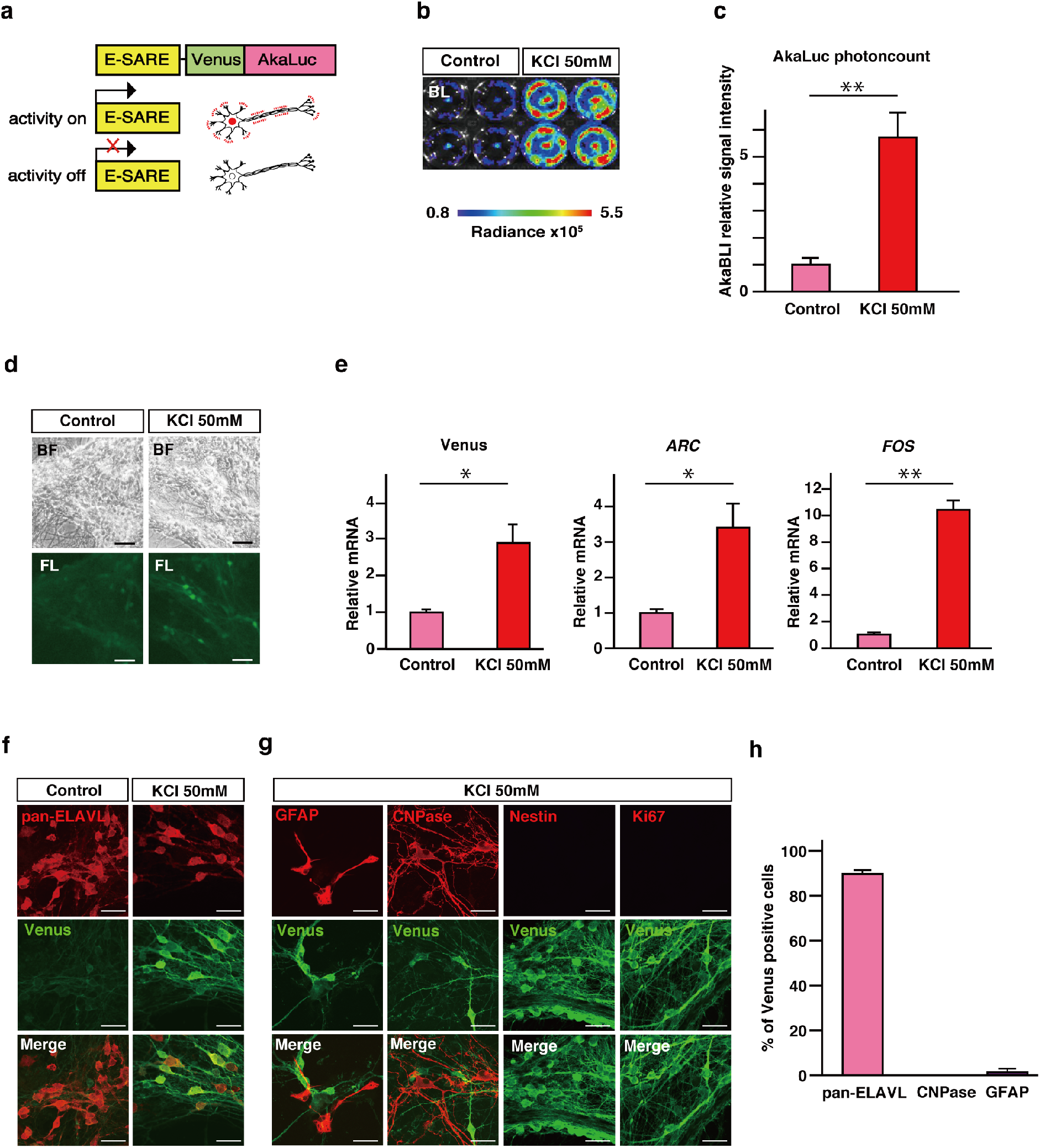
BLI of NS/PC-derived cells stimulated with high potassium *in vitro*. **(a)** Schematic illustration of the E-SARE-Venus-AkaLuc (ESAL) construct, which was used to expresses the Venus-fused AkaLuc luminescent enzyme under control of the promoter E-SARE. When a neuron was activated, the promoter E-SARE drove high expression of the downstream reporter gene Venus-AkaLuc. **(b)** Comparative BLI of *in vitro*-cultured cells stimulated with 50mM KCl for six hours on the right (n = 4) or without stimulation (n = 4) on the left. The colour of the bars indicates the total bioluminescence radiance (photons/sec/cm^2^/str). **(c)** Quantitative analyses of the relative luminescence intensity of ESAL-NS/PC-derived cells with or without the addition of 50 mM KCl *in vitro* (n = 4 each). **(d)** Microscopic bright-field image and GFP fluorescence image of ESAL-NS/PC-derived cells with or without the addition of 50 mM KCl *in vitro.* Scale bar, 50 µm. **(e)** The results of quantitative real-time PCR analyses of the gene expression of Venus, *ARC*, and *FOS* in cells within the same well as shown above (n = 4 each). **(f), (g)** Representative images of Venus-expressing grafted cells stained with pan-ELAVL: neurons, with or without the addition of 50 mM KCl (**f**), or stained with human GFAP; astrocytes, CNPase; oligodendrocytes, Nestin and Ki-67; immature cells, with the addition of 50 mM KCl (**g**). Scale bar, 20 µm. **(h)** Percentage of cells positive for cell type-specific markers among the Venus+ cells stimulated with 50 mM KCl. Values are the mean ± SEM; *p < 0.05, **p < 0.01. Statistical analyses were performed using the two-sided unpaired Student’s *t* test in c and e. Individual t-values and degrees of freedom: **c;** t (6) = 11.59, p = 2.5×10^-5^. **e;** Venus t (6) = 3.034, p = 0.023. *Arc* t (6) = 2.705, p = 0.035. *Fos* t (6) = 7.057, p = 4.1×10^-4^.

To profile the temporal resolution of the ESAL system, we examined the time course of bioluminescence after a brief bout of neuronal stimulation (4AP+BIC) that facilitates action potential firing and potentiates glutamatergic transmission (Figure 1–figure supplement 1a). Neuronal activity-dependent bioluminescence was detected approximately 4 hours after stimulation, reached its peak at 6 hours, and returned to the basal level at 24 hours (Figure 1–figure supplement 1b). These results indicate that an increase in bioluminescence in the ESAL system reflects cumulative neuronal activity that persisted during a period ranging from 4 to10 hours prior to measured BLI.

### Monitoring the activity of neuronal grafts in an SCI model by ESAL

Next, NOD/ShiJic-scidJcl (NOD-SCID) mice were subjected to C4 spinal cord transection, and 9 days after injury, ESAL-NS/PCs were transplanted into the injury sites (Figure 2a). The grip strength test revealed that ESAL-NS/PC transplantation (TP), but not the PBS injection (PBS), induced a significantly better recovery of motor function (Figure 2b).

**Figure 2.**
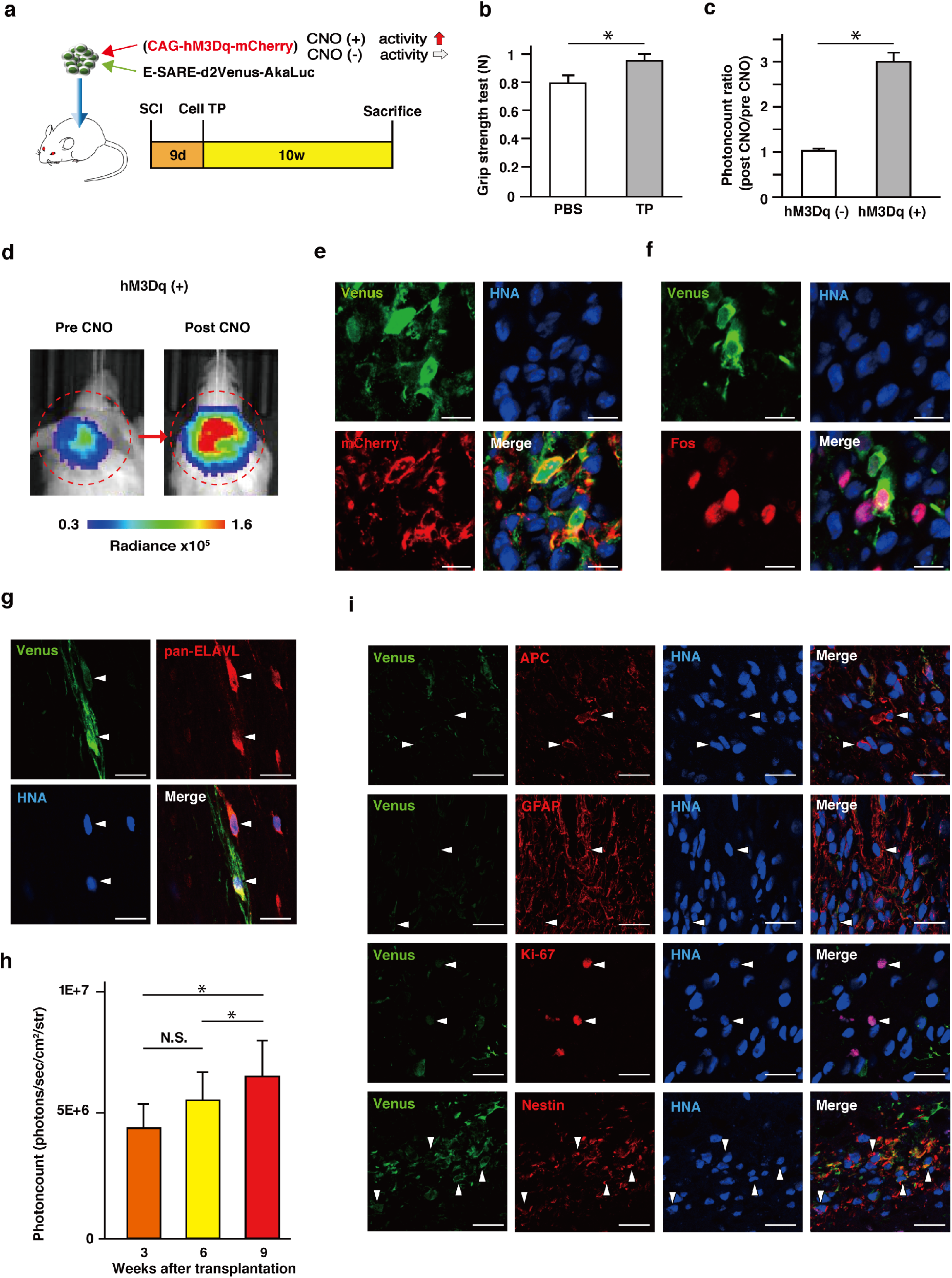
*In vivo* application of ESAL-expressing NS/PC transplantation after cervical spinal cord injury. **(a)** Schematic illustration of *in vivo* experiments. Transplantation of co-infected NS/PCs (positive control mice; CAG-hM3Dq-mCherry, which contains a hM3Dq and mCherry fusion protein, was transduced to ESAL NS/PCs via lentivirus as well) (**d**–**f**), or transplantation of ESAL-expressing NS/PCs (**g**–**i**) was performed 9 days after C4 dorsal hemisection. **(b)** Grip strength testing of the PBS group and TP group was performed before sacrifice 70 days after TP (TP group: n = 7, PBS group: n = 6). Values are the mean ± SEM; *p < 0.05, **p < 0.01. Statistical analyses were performed using the two-sided unpaired Student’s *t* test. Individual t-values and degrees of freedom: t (11) = 2.285, p = 0.043. **(c)** The ratios of luminescence intensity (post CNO/pre CNO) of ESAL-expressing NS/PC-transplanted mice (hM3Dq minus, n = 4) and the positive control mice (hM3Dq plus, n = 3) are shown. Statistical analyses were performed using the two-sided unpaired Student’s *t* test. Individual t-values and degrees of freedom: t (5) = 3.977, p = 0.011. **(d)** Representative IVIS images of a positive control mouse (pre and post CNO). The circle shows the region of interest (ROI) in the cervical spine. The colour of the bars indicates the total bioluminescence radiance (photons/sec/cm^2^/str). **(e), (f)** Immunohistological images of a positive control mouse 6 weeks after transplantation; labelled with Venus (green), mCherry (red), and HNA (human cells) (blue) (**e**) or labelled with Venus (green), Fos (red), and HNA (blue) (**f**). Scale bars, 10 µm. **(g)** A representative image of an ESAL-expressing NS/PC-transplanted mouse labelled with Venus (green), pan-ELAVL (red), and HNA (blue) (arrowheads). Scale bars, 20 µm. **(h)** Time-dependent change in graft luminescence intensity of ESAL-expressing NS/PC-transplanted mice at 3, 6, and 9 weeks after transplantation (n = 12 mice). Values are the mean ± SEM; *p < 0.05, **p < 0.01. Statistical analyses were performed using the Friedman test. Individual p values: three and six weeks; 0.082, six and nine weeks; 0.016, three and nine weeks; 0.038, Fisher’s LSD. **(i)** A representative image of an ESAL-expressing NS/PC-transplanted mouse labelled with Venus (green); APC (oligodendrocyte), GFAP (astrocyte), and Ki-67/Nestin (red); and HNA (blue) (arrowheads). Scale bars, 20 µm.

To artificially manipulate the neuronal activity in the grafted cells, we introduced hM3Dq, a stimulatory chemogenetic receptor and designer receptor exclusively activated by designer drugs (DREADDs), which enables the graft to be activated upon administration of its ligand (Nichols & Roth, 2009; Roth, 2016) into ESAL-NS/PCs. When the grafts were activated by administration of the hM3Dq ligand clozapine N-oxide (CNO), bioluminescence was significantly upregulated (Figure 2c, d). Consistently, in immunohistochemical analyses, Venus protein expression was detected exclusively in hM3Dq-expressing (mCherry+) graft cells (95.3 ± 2.8%) (Figure 2e). In addition, Venus+ cells were mostly immunopositive for Fos, a marker and an IEG (68.9 ± 3.0%) (Figure 2f). These data suggested that the ESAL system successfully labelled active ensembles in neurons grafted into the SCI model mice.

Even without artificial activation by hM3Dq, we found Venus expression in a portion of neuronal grafts (Figure 2g), suggesting that the ESAL system may report neurons showing spontaneous activity in the graft. This is in line with the time course of ESAL bioluminescence elevation measured chronologically after ESAL-NS/PC transplantation, suggesting that a progression of neuronal differentiation and maturation of NS/PCs precedes a significant increase in bioluminescence detection (Figure 2h). In keeping with this idea, we confirmed only a few immature non-neural cells as the origin of the Venus+ cells (Figure 2i).

### Graft neurons integrates into the host nervous system

Previous reports have suggested that host neural circuits gain projection to the graft, whose activity was regulated and linked to the host neuronal dynamics (Kadoya et al., 2016). Using the ESAL system, we examined the effect of host activity on neuronal graft activity at the individual and circuit level. First, we monitored ESAL bioluminescence over the day and found that ESAL bioluminescence was regulated highest around noon and lowest at night (Figure 3–figure supplement 1a). Given that ESAL bioluminescence reflects cumulative neuronal activity approximately 6 hours before observation (Figure 1–figure supplement 1a, b), this result indicates that graft neuronal activity has diurnal variations consistent with the peak and the trough activity of the host animals during the nocturnal and day periods, respectively. To further determine how much host activity influences graft activity at the individual level, we next utilized long-term anaesthesia (a combination of midazolam, medetomidine hydrochloride, and butorphanol) to mimic sleep (Figure 3a). The respiration rate of the mice suggested that the anaesthesia was active for at least 6 hours following administration. To increase the anaesthetic time to 9 hours, we additionally administered isoflurane for the remaining 3 hours. The ESAL bioluminescence decreased to nearly half of the initial level after long-term anaesthesia (Figure 3b, c). This decrease was significant at both 6 and 9 weeks post-transplantation (Figure 3c). Additionally, we sought to confirm that three types of mixed anaesthetic agents cannot directly alter the activity of graft neurons, using *in vitro*-cultured neurons. Indeed, BLI signal intensity was not notably reduced by the anaesthetic agents (Figure 3d, e). Taken together, these data imply that graft neuronal activity is associated with the daily activities of hosts on an individual level.

**Figure 3.**
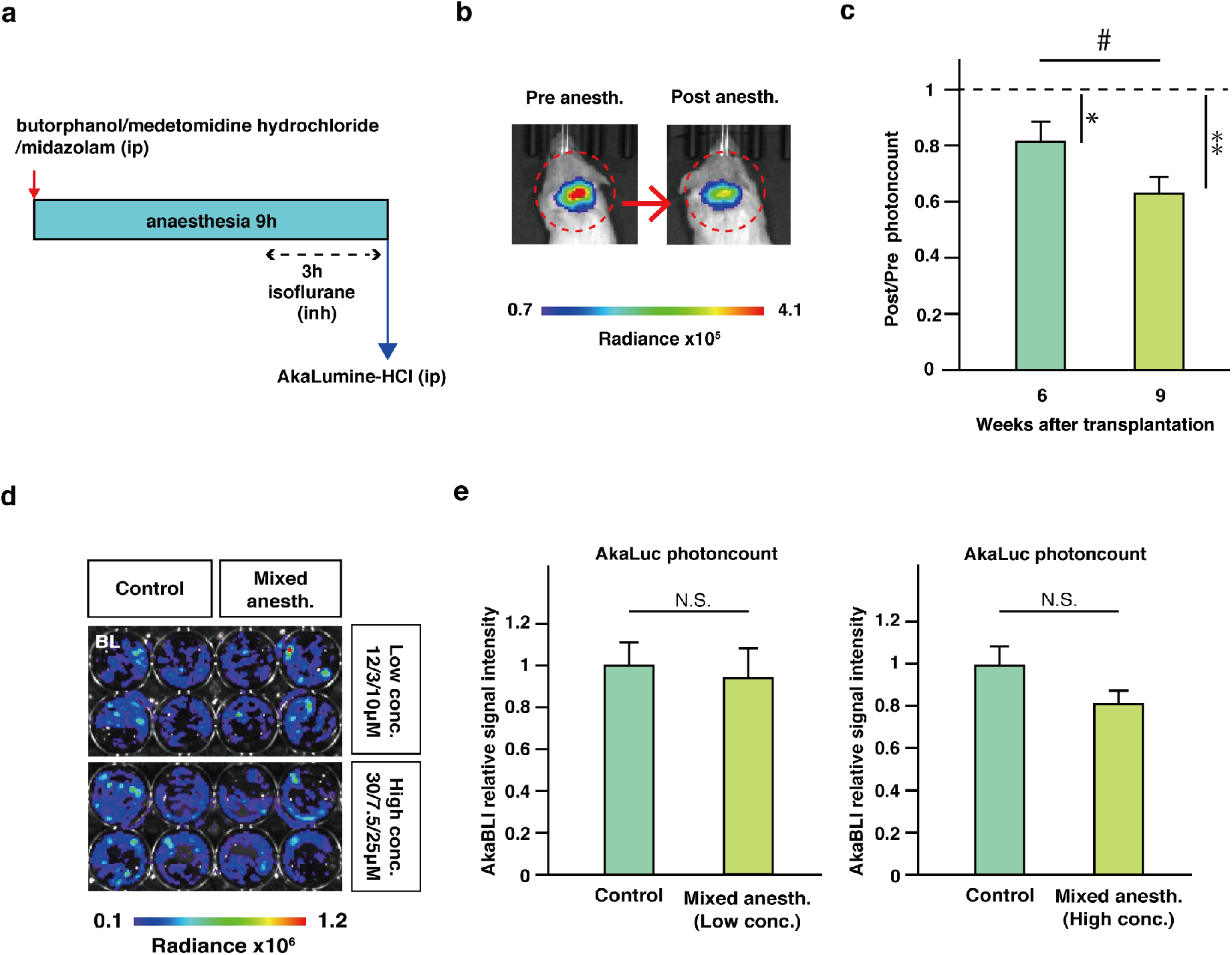
Continuous anaesthetization of ESAL-expressing NS/PC-transplanted mice. **(a)** Schematic illustration of long-term continuous anaesthesia; anaesthesia with a mixture of three types of anaesthetic agents (butorphanol, medetomidine hydrochloride, midazolam), followed by inhalation anaesthesia for up to nine hours. **(b)** Representative IVIS images of an ESAL-expressing NS/PC-transplanted mouse before and after long-term continuous anaesthesia 70 days after transplantation. The circle shows the region of interest (ROI) in the cervical spine. The colour of the bars indicates the total bioluminescence radiance (photons/sec/cm^2^/sr). **(c)** IVIS photon count ratio before and after continuous anaesthesia at six and nine weeks after transplantation (n = 5 mice). **(d)** Comparative BLI of *in vitro*-cultured cells with or without the addition of the mixed anaesthetic agents at two different concentrations for six hours (n = 4, 4 each). The final concentration of each anaesthetic agents is written on the right hand. The colour of the bars indicates the total bioluminescence radiance (photons/sec/cm^2^/str). **(e)** Quantitative analyses of the relative luminescence intensity of ESAL-NS/PC-derived cells with or without the addition of the mixed anaesthetic agents *in vitro* (n = 4, 4 each). Values are the mean ± SEM; *p < 0.05, **p < 0.01, ^#^p < 0.05. Statistical analyses were performed using the two-sided paired Student’s *t* test in **c**, and the two-sided unpaired Student’s *t* test in **e**. Individual t-values and degrees of freedom: **c;** 6 and 9 weeks; t (8) = 2.754, p = 0.025, 6 weeks; t (8) = 2.502, p = 0.037, 9 weeks; t (8) = 4.456, p = 2.1×10^-3^. **e;** left: t (6) = 0.342, p = 0.744, right: t (6) = 1.821, p = 0.118.

### Host neural circuit inputs regulate graft activity

We next investigated whether and to what extent host circuit-level activity directly regulates graft neuronal activity. We focused on the corticospinal tract (CST), one of the main descending circuits that plays pivotal roles in sensorimotor control, and artificially manipulated CST activity by injection of the motor cortex with adeno-associated virus (AAV) encoding hM3Dq-mCherry under control of the human synapsin I promoter (Figure 4a). Three to four weeks after AAV injection, we confirmed that hM3Dq-mCherry permitted efficient anterograde labelling to the C4 lesion site through the CST (Figure 4b). This selective labelling of the CST projections to the lesion site suggested that CST fibres innervated the graft (Figure 4c). Indeed, we found that host-to-graft synapses had formed (Figure 4d), indicative of CST-driven control of the graft. To test this directly, the CST was activated by the administration of the hM3Dq ligand CNO, and we detected an activity-induced increase in Venus expression fused with AkaLuc in graft cells by immunohistochemical analysis (Figure 4e, f). Furthermore, the increase in graft activity induced by artificial CST stimulation upon CNO treatment was also confirmed *in vivo* by BLI measurements through an increase in ESAL photon counts of the graft (Figure 4g, h). These results suggest that host CST inputs innervate and regulate graft activity.

**Figure 4.**
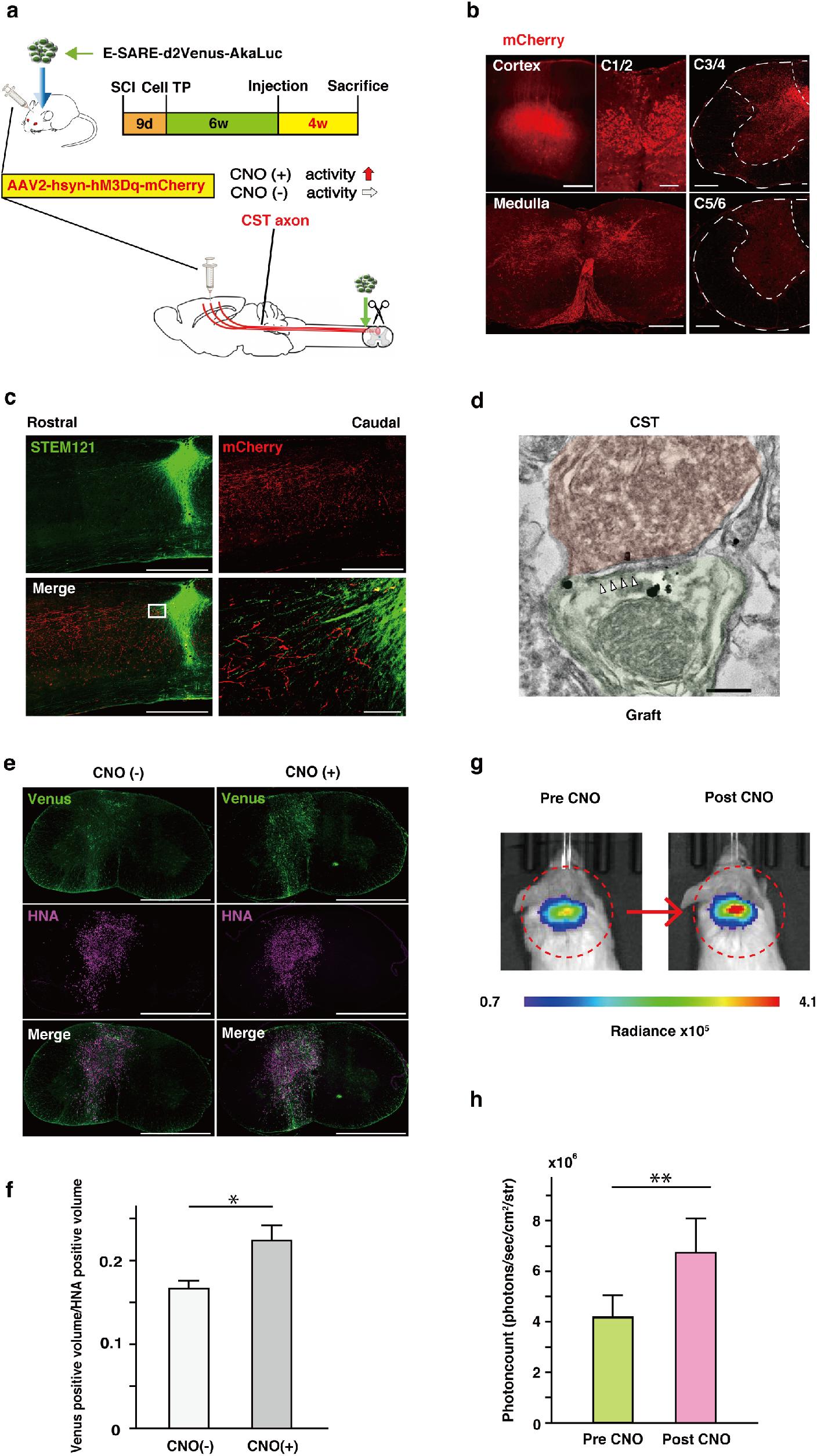
CST stimulation via clozapine N-oxide (CNO) administration through the DREADD system in ESAL-NS/PC-transplanted mice. **(a)** Schematic illustration representing the time schedule of the *in vivo* experiments. Six weeks after transplantation of ESAL-expressing NS/PCs, all mice were subjected to AAV injection into motor cortex. Ten weeks after transplantation, luminescence measurements were performed before sacrifice. **(b)** Confirmation of mCherry-labelled cell bodies in the cerebral cortex (scale bars, 250 µm), mCherry-labelled axons in the medulla (scale bars, 500 µm), and cervical spinal cord 1/2 level (scale bars, 100 µm). mCherry+ CST axon ends were observed in the grey matter rostral to the C4 lesional area (scale bars, 250 µm), and few CST axons were observed caudal to the lesion (scale bars, 250 µm). **(c)** Representative sagittal images 400 µm from the midsagittal plane around the injured spinal cord. Transplanted cells were stained with STEM121 (human cytoplasmic marker), and the CST was labelled with mCherry. Scale bars, 1000 µm, 50 µm. **(d)** Double immunoelectron microscopic image of the synaptic connection between an mCherry+ CST neuron with DAB staining and a Venus+ graft neuron with Fluorogold staining. Anti-mCherry labelling was localized at the membrane of the CST neuron, and Venus proteins were detectable as black dots, mainly in the cytoplasmic membrane. Arrowheads, post synaptic density. Scale bar, 500 nm. **(e)** Representative axial cervical spinal cord images of Venus and HNA staining; tissue from mice with or without CST activation by 5 mg/kg CNO administration 70 days after transplantation is shown. Scale bar, 1000 µm. Venus is a cytoplasmic protein, while HNA is an anti-nuclear antigen. **(f)** Comparison of calculated Venus+ volume/HNA+ volume; group sacrificed with CST activation (n = 4 mice) and group sacrificed without CST activation (n = 4 mice). Anti-GFP antibody is used to label Venus protein. **(g)** Representative IVIS images of an ESAL-expressing NS/PC-transplanted mouse before and after CST activation (the same individual shown in Figure 3b, 4e[right]). The circle shows the region of interest (ROI) in the cervical spine. The colour of the bars indicates the total bioluminescence radiance (photons/sec/cm^2^/str). **(h)** Comparison of luminescence intensity for ESAL-expressing NS/PC-transplanted mice with or without CST activation for each mouse 70 days after transplantation. (n=10 mice). Values are the mean ± SEM; *p < 0.05, **p < 0.01. Statistical analyses were performed using the two-sided unpaired Student’s *t* test in **f** and the two-sided paired Student’s *t* test in **h**. Individual t-values and degrees of freedom: **f;** t (6) = 2.573, p = 0.042. **h;** t (18) = 2.963, p = 8.3×10^-3^.

## Discussion

Here, we developed a novel bioimaging system, ESAL, by combining a sensitive and accurate red-shifted bioluminescence, AkaBLI, and the neuronal activity-dependent promoter E-SARE. This ESAL system efficiently labels active neurons in neural grafts in the injured spinal cord. By using this non-invasive ESAL imaging system, we have demonstrated the direct association between graft activity and the host circuit/behaviour-level activity.

The results of this study demonstrate that the ESAL system is a non-invasive method for *in vivo* imaging of graft activity in the injured spinal cord. The ESAL system can image spontaneous neuronal activity and reveal the interaction between the graft and host neural circuits. Although some groups have reported host-graft connectivity in SCI models by using electrophysiological (Kadoya et al., 2016) or calcium imaging techniques (Ceto, Sekiguchi, Takashima, Nimmerjahn, & Tuszynski, 2020), these experiments were not performed in the living SCI animals. In contrast, it is noted that the ESAL system can be used in an entirely non-invasive manner, and we showed *in vivo* that host activity at the individual level, such as sleep and long-term anaesthesia, directly influences graft activity.

Neuronal relay is a core mechanism through which NS/PC transplantation can be used to treat SCI (Ceto et al., 2020; Dell’Anno et al., 2018; Kadoya et al., 2016). Relay circuits can be established between host descending axons and newly differentiated neurons from transplanted NS/PCs. However, the lack of non-invasive *in vivo* measurement methods has precluded our understanding on how the neural grafts functionally integrate into the host circuits, and how the graft activity contributes to the behavioural recovery. Our ESAL system can monitor graft activity in the context of various host behaviours, leading to the mechanistic elucidation of neuronal relay formation and its contribution to functional recovery.

The bioluminescence intensity of NS/PC-derived cells increased continuously until 9 weeks after transplantation, which suggests that spontaneous graft activity increased concomitantly. This is in keeping with a previous study from our group suggesting that more than 6 weeks is required for neuronal maturation in grafts (Nori, Nakamura, & Okano, 2017). Several reports using human iPSC-derived brain organoids also indicated that synapse formation and spontaneous activity begin at 4–6 weeks (Wilson, 2018). Based on their findings and ours, the ESAL system results could be reflective of neuronal maturation and synaptic formation *in vivo*.

The debate remains controversial on which subtypes of NS/PCs with different regional identities are the most appropriate for cell therapy (Kadoya et al., 2016; Kajikawa et al., 2020; Watanabe et al., 2004). The ESAL system will enable determining the quality and suitability of various NS/PC subtypes integrated to the host. How the graft contributes to further tissue regeneration and motor function restoration has remained elusive. Previous reports showed that pretreatment with a γ-secretase inhibitor (GSI) promoted the maturation of NS/PC-derived neurons by inhibiting Notch signalling (Okubo et al., 2016); furthermore, Nogo receptor antagonists facilitated raphespinal tract regeneration (Ito et al., 2018), and some synapse organizers were suggested to accelerate synaptic formation (Suzuki et al., 2020). This system will also be valuable in evaluating the effect of these molecular components on NS/PC grafts and the host circuitry around the injured spinal cord.

Thus, this ESAL system has the potential to reveal synaptic input contribution from each descending pathway to graft neurons in motor function recovery. Functions of the CST include the control of afferent inputs, spinal reflexes and motor neuron activity (Lemon & Griffiths, 2005). Furthermore, the CST is generally recognized as the principal motor pathway for voluntary movements in humans (Welniarz, Dusart, & Roze, 2017). As in rodents, the CST commands only gripping ability involving digital flexors, which largely depends on the dorsal and dorsolateral CST, as well as reaching tasks (Schrimsher & Reier, 1993). According to previous studies, the reticulospinal tract may be more important than the CST for motor function in rodents (Ballermann & Fouad, 2006; Lemon, 2008). It would be interesting to verify the transplantation site by using this system to enhance the effect of cell therapy.

It is important to note that ESAL system does not monitor real-time ongoing neuronal activity, but reflects cumulative activity-dependent gene expression that integrates activity in the recent past due to the prolonged E-SARE promoter activity after neuronal stimulation. One solution to improve temporal resolution is to utilize an indicator of real-time neuronal activity, such as the intracellular calcium concentration. Indeed, a calcium-dependent luciferase system, Orange CaMBI, has been reported (Oh et al., 2019). However, this system is based on NanoLuc-furimazine and might not be appropriate for neural grafts due to its low substrate permeability through the blood-brain barrier (Edinger et al., 1999; Su et al., 2020). Another possible solution is the use of another activity-dependent promoter, such as the *Fos* promoter (Schilling, Luk, Morgan, & Curran, 1991), which is a commonly used activity-dependent promoter that is regulated at a very low level because of the autorepression of *Fos* transcription by the Fos protein (Lucibello, Lowag, Neuberg, & Müller, 1989; Morgan & Curran, 1991). Thus, to amplify *Fos* promoter-dependent expression, a tet-inducible system is usually added due to transcriptional activation by tetracycline (Iwano et al., 2018). However, this *Fos-*tet double-reporter system requires substantial time for gene expression. In contrast, the ESAL system drives reporter expression at a high enough level to be used alone for graft monitoring, achieving gene expression over a relatively short period.

Although we have mainly focused on the host-to-graft connection between the host descending neurons and the graft, further studies are needed to assess the graft-to-host interaction between the graft and the host spinal neurons, such as motor neurons. The ESAL system is available for not only grafts but also host neurons through the use of AAV, because ESAL can be packaged into a single AAV due to its small size. For example, it would be feasible to detect the activity of spinal motor neurons by transfecting an AAV retrograde viral vector into the neuromuscular junction (Tervo et al., 2016). Future studies will extend our understanding of the mutual interaction between host and graft.

In conclusion, this study introduces a new *in vivo* system to monitor grafted cell-derived neurons and provides important information on the importance of the link between graft activity with the host neuronal circuit and behaviour. We demonstrated *in vivo* that host-to-graft synaptic connectivity was functionally established after NS/PC transplantation for SCI. One way to improve cell therapy would be to further enhance this connectivity. Promoting neuronal maturation or synaptic formation might render cell therapy with this system more effective in the future.

## Materials and Methods

### Lentiviral vector construction

To construct a lentiviral vector for ESAL, the promoter E-SARE and Venus-AkaLuc-PEST cDNA were cloned into the lentiviral vector CSII. To construct a lentiviral vector for ubiquitous DREADD activation, hM3Dq-mCherry cDNA was polymerase chain reaction (PCR)-amplified from pAAV-hSyn-hM3D(Gq)-mCherry (Addgene plasmid #50474) and transferred to the lentiviral vector CSIV with the CAG promoter, which is a hybrid construct consisting of the cytomegalovirus (CMV) enhancer fused with the chicken beta-actin promoter (Sakai, Mitani, & Miyazaki, 1995).

### Lentiviral vector preparation

Recombinant lentiviral vectors were produced by transient transfection of three plasmids into HEK 293T cells, pCAG-HIVgp, pCMV-VSV-G-RSV-Rev, and the lentiviral vector (CSII-E-SARE-Venus-AkaLuc-PEST or CSIV-CAG-hM3Dq-mCherry), as previously described (Iida et al., 2017; Kojima et al., 2019; Miyoshi, Blömer, Takahashi, Gage, & Verma, 1998).

### NS/PC culture and lentiviral transduction

NS/PCs were generated from the human iPSC line 414C2 (Okita et al., 2011) by previously described methods (Nori et al., 2017; Okada et al., 2008; Okubo et al., 2016). Briefly, embryoid bodies (EBs) were generated from iPSCs grown in suspension for 30 days. The EBs were then dissociated into single cells using TrypLE Select (Thermo Fisher Scientific, MA, USA) and cultured in suspension in KBM neural stem cell medium (Kohjin Bio, Saitama, Japan) supplemented with B-27 (Thermo Fisher Scientific), 20 ng/ml FGF-2 (PeproTech, NJ, USA), and 10 ng/ml human leukaemia inhibitory factor (hLIF; Merck KGaA, Hesse, Germany) for 12 days. These primary neurospheres were passaged every 10–14 days. Lentiviral infection was performed after the first passage, and tertiary neurospheres were used for the following experiments. For the transplantation experiments, GSI treatment was applied on the day before transplantation as described previously (Okubo et al., 2016).

### *In vitro* neuronal differentiation

Dissociated tertiary neurospheres were plated onto 24-well plates precoated with mouse astrocytes at a density of 1×10^5^ cells/well. The mouse astrocytes had been extracted from the E17 mouse cerebral cortex (5×10^4^ cells/well). Cells were cultured at 37 °C in 5% CO_2_ and 95% air for 50 days in neuronal maturation medium consisting of Neurobasal Plus Medium (Thermo Fisher Scientific) supplemented with B-27 Plus Supplement (Thermo Fisher Scientific), GlutaMAX (Thermo Fisher Scientific), Culture One Supplement (Thermo Fisher Scientific), and L-ascorbic acid (200 µM) (Sigma-Aldrich, MO, USA). For neuronal stimulation, potassium chloride was added at a concentration of 50 mM and incubated for six hours. A three-drugs mixture (butorphanol; Vetorphale, Meiji Seika Pharma Co., Ltd., Tokyo, Japan/ medetomidine hydrochloride; Domitor, Nippon Zenyaku Kogyo Co., Ltd., Fukushima, Japan/midazolam; Midazolam, Sandoz K.K., Tokyo, Japan) was added at two different concentrations; 12/3/10 µM (low concentration), 30/7.5/25 µM (high concentration) (Hsiao et al., 2004; Wang et al., 2018).

### SCI modelling and transplantation

Eight-week-old female NOD-SCID mice (20–22 g; Oriental Yeast Co., Ltd., Tokyo, Japan) were anaesthetized by intraperitoneal injections of ketamine (60 mg/kg) and xylazine (10 mg/kg). The laminal arch of the vertebrae at the C4 level was removed, and the dorsal surface of the dura mater was exposed. A tungsten wire knife (McHugh Milieux, David Kopf Instruments, CA, USA) was inserted 0.6 mm from the dorsal surface and raised 0.5 mm to transect the dorsal column as described previously with slight modifications (Kadoya et al., 2016). Nine days after the injury was made, 5×10^5^ NS/PCs per 2 µl were transplanted into the lesion area at a rate of 1 µl/minute using a metal needle with a 10-µl Hamilton syringe and a stereotaxic microinjector (KDS 310; Muromachi Kikai, Tokyo, Japan). An equal volume of PBS was injected instead into control mice. After SCI and transplantation, 12.5 mg/kg ampicillin was administered intramuscularly. To determine the therapeutic effect of transplantation on motor function, all animals within an experimental group that underwent C4-CST lesions were randomly assigned to either TP or PBS group. All animal experiments were approved by the Ethics Committee of Keio University and performed in accordance with the Guide for the Care and Use of Laboratory Animals (National Institutes of Health, MD, USA).

### Behavioural analysis

Recovery of motor function following cell transplantation or PBS injection was assessed using the grip strength test before sacrifice (cell transplantation group; n = 7, PBS group; n = 6). The trial consisted of five separate pulls. The highest and lowest forces were excluded, and the remaining three forces were averaged (Forgione, Chamankhah, & Fehlings, 2017). The grip strength test was performed using a digital force gauge (Shimpo, Kyoto, Japan) and wire mesh attachment device (Muromachi Kikai).

### Anterograde labelling and activation of the CST

Six weeks after transplantation, AAV2-hsyn-hM3Dq-mCherry (Addgene #50474-AAV2; 7.38 ×10^12^ vg/ml) was injected into the bilateral sensorimotor cortex at four sites (500 nL/point; coordinates = 1 mm rostral and 1.4 mm lateral to bregma, 1 mm posterior and 1 mm lateral to bregma; depth = 0.7 mm) at a rate of 100 nL/minute through a pulled glass micropipette (calibrated micropipette, 1–5 µL; Funakoshi, Tokyo, Japan). Nine to 10 weeks after transplantation, CNO (Enzo Life Sciences, NY, USA) was intraperitonially administered at a concentration of 5 mg/kg. To investigate whether CST activation alter the activity of neural grafts, 15 subjects underwent C4-CST lesions, TP, and anterograde labelling of CST axons. To be included in analysis, the graft should be clearly detected throughout the experiments (3, 6, 9 weeks after transplantation), and the CST axons should be successfully transected and labelled. Out of the 15 animals, three animals were excluded because of death before final measurement, and two animals were excluded because of poor labelling of CST axons.

### Luminescence measurement

Bioluminescence images were acquired using the IVIS Spectrum system (Perkin Elmer, MA, USA). For *in vitro*-cultured neurons, bioluminescence was measured immediately after treatment with 300 µM AkaLumine-HCl (FUJIFILM Wako Pure Chemical, Osaka, Japan). Animals were imaged on a schedule of three, six, and nine weeks after transplantation under inhalation anaesthesia (2% isoflurane and oxygen) or under three types of mixed anaesthesia (5 mg/kg butorphanol, 0.75 mg/kg medetomidine hydrochloride, and 4 mg/kg midazolam) (Kawai, Takagi, Kaneko, & Kurosawa, 2011) for nine hours, followed by inhalation of 2% isoflurane and oxygen. The signal was measured for 15 minutes after 50 µl of AkaLumine-HCl (60 mM) and saline solution had been intraperitoneally injected. The region of interest (ROI) was set immediately above the cervical cord, and the peak intensity, observed at approximately 10 minutes in most cases, was recorded. The measurement parameters were as follows: *in vitro*; exposure time = 1 s, binning = 8, field of view = 13.4 cm, and f/stop = 1; *in vivo*; exposure time = 60 s, binning = 8, field of view = 23 cm, and f/stop = 1. All the images were processed with Living Image software (IVIS Imaging Systems), and the signal intensity is expressed as the photon count in units of photons/sec/cm^2^/str. Each result was displayed as a pseudo-coloured photon count image superimposed on a grey-scale anatomic image.

### *In vitro* luminescence measurement in mouse neurons over time

Hippocampal neurons isolated from E16 mouse embryos were cultured on 24-well plates coated with fibronectin (Sigma-Aldrich). The cells were infected with lentivirus-E-SARE-Venus-AkaLuc-PEST on DIV5, silenced with TTX (1 µM, Tocris, Bristol, UK) on DIV7 and then stimulated with 4AP (250 µM, Tocris) and bicuculline (50 µM, Sigma-Aldrich) in the absence of TTX for 10 min on DIV8. After stimulation, the neurons were silenced again with medium containing 1 µM TTX. At designated time points (0, 2, 4, 6, 8, 10, and 24 hours after brief stimulation), bioluminescence images were acquired using the IVIS Spectrum system immediately after treatment with 300 µM AkaLumine-HCl.

### qPCR

Total RNA was extracted by using an RNeasy Micro Kit (Qiagen, Inc., Hilgen, Germany), and cDNA was synthesized by reverse transcription with ReverTra Ace qPCR RT master mix (Toyobo Co., Ltd., Life Science Department, Osaka, Japan). Quantitative PCR (qPCR) was performed using Step One Plus (Applied Biosystems, CA, USA) following the manufacturer’s instructions. The expression levels of each gene were normalized to that of *ACTB* using the comparative ΔΔCT method. We used the following manufactured primers (Thermo Fisher Scientific) against human DNA sequences: *FOS* (Hs01119266_g1), *ARC* (Hs01045540_g1), and *ACTB* (Hs03023943_g1). Additionally, *EGFP* (Mr00660654_cn) was used to detect Venus expression.

### Immunostaining

*In vitro*-cultured cells were fixed with 4% paraformaldehyde (PFA) for 15 minutes. All mice were deeply anaesthetized and transcardially perfused with 4% PFA 10 weeks after injury. The dissected spinal cords were embedded in optimal cutting temperature compound (Sakura Finetek, Tokyo, Japan) and sectioned in the axial plane at a thickness of 12 µm on a cryostat (Leica Biosystems, Wetzlar, Germany). The samples were stained with the following primary antibodies: anti-GFP (goat IgG, 1:500, Rockland, PA, USA), anti-mCherry (rabbit IgG, 1:400, Abcam, Cambridge, UK), anti-pan-ELAVL (mouse IgG1, 1:200, Sigma-Aldrich), anti-GFAP (rabbit IgG, 1:2000, Proteintech, IL, USA), anti-APC (mouse IgG2b, 1:300, Abcam), anti-human GFAP (mouse IgG1, 1:2000, Takara Bio, Shiga, Japan), anti-CNPase (mouse IgG1, 1:2000, Sigma-Aldrich), anti-Ki-67 (rabbit IgG, 1:2000, Leica Biosystems), anti-Nestin (rabbit IgG, 1;200, IBL, Gunma, Japan), anti-HNA (msIgG1, 1:100, Millipore, Darmstadt, Germany), anti-Fos (rabbit IgG, 1:400, Abcam), and STEM121 (msIgG1, 1:200, Takara Bio). The nuclei were stained with Hoechst 33258 (10 µg/ml, Sigma-Aldrich). All images were obtained using a fluorescence microscope (BZ-X710; Keyence, Osaka, Japan) or confocal laser scanning microscope (LSM 780; Carl Zeiss, Jena, Germany).

### Quantitative analysis of the tissue sections

Quantitative analysis of the tissue sections following SCI and transplantation was performed as described previously (Kajikawa et al., 2020). Three-dimensional analysis of the Venus+ volume (graft activity)/HNA+ volume (human cells) was performed as follows. Axial sections were prepared from eight animals, and the Venus+ area and HNA+ area were determined using ImageJ; the volume was then calculated as follows:

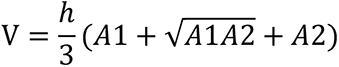

where A1 and A2 are the areas of two consecutive sections, and h is the distance between them (480 µm).

### Double immunoelectron microscopy

The detailed procedure used for pre-embedding immunoelectron microscopic analysis has been described previously (Shibata et al., 2019). Briefly, frozen spinal cord sections on glass slides were thawed, dried and autoclaved in citrate acid buffer (pH 6.0), followed by blocking treatment [5.0% Block Ace (DS Pharma Biomedical, Osaka, Japan) solution with 0.01% saponin in 0.1 M PB]. The samples were stained with the primary antibodies anti-GFP (goat IgG, 1:100, Rockland), anti-mCherry (rabbit IgG, 1:100, Abcam) and the secondary antibodies anti-rabbit biotin (donkey IgG, 1:800, Jackson ImmunoResearch, PA, USA) and Alexa Fluor 488 Nanogold-conjugated rabbit anti-goat IgG antibody (1:100, Nanoprobes, NY, USA) before staining with Hoechst 33258. We used the following supplements: the ABC complex (Vectastain Elite ABC Kit; Vector, CA, USA), TSA Plus biotin (NEL749A001KT; PerkinElmer, MA, USA), SA-Alexa Fluor 555 (1:1000, Thermo Fisher Scientific), SA-HRP (1:100, Vector), and 3,3’-diaminobenzidine (DAB) tablets (FUJIFILM Wako Pure Chemical). Ultrathin sections (80-nm thickness) were prepared with a diamond knife, collected on copper mesh grids (#100 or #150 Veco, Nisshin EM, Tokyo, Japan), and stained with uranyl acetate and lead citrate in plastic tubes for 10 minutes each. The sections were examined with a transmission electron microscope (TEM, JEM-1400Plus, JEOL, Tokyo, Japan) at 100 keV.

### Statistical analysis

For comparisons between two groups, a two-tailed Student’s t test was used. The Friedman test followed by a post hoc Fisher’s test was used for analysis of the data in Fig. 2h. For all statistical analyses, differences were considered significant at p < 0.05. All data are presented as the mean ± SEM. IBM SPSS Statistics (ver. 26) was used for all calculations.

### Data availability

Source data are provided with this paper. The data that support the findings are available on request from the corresponding authors.

## Acknowledgements

We thank S. Yamanaka at CiRA (Kyoto University) for supplying the 414C2 human iPSCs. We thank K. Tanaka at the Department of Psychiatry (Keio University) for technical and conceptual guidance. We thank H. J. Okano and M. Hasegawa at the Division of Regenerative Medicine (Jikei University) for their assistance with the experiments. We are grateful for the assistance of H. Miyoshi, S. Nori, O. Tsuji, S. Ito, Y. Hoshino, Y. Tanimoto, T. Shibata, S. Hashimoto, Y. Suematsu, Y. Saijyo, T. Nishijima, T. Tanaka, K. Ito, L. Tao, and K. Nakanishi, who are all members of the spinal cord research team at the Department of Orthopaedic Surgery and Physiology (Keio University). We also thank T. Harada, K. Yasutake, and M. Akizawa for their assistance with the experiments and animal care.

## Funding

This work was supported by the Japan Agency for Medical Research and Development (AMED) (grant no. JP20bm0204001, JP19bm0204001, JP20bk0104017, and JP19bk0104017 to H.O. and M.N.) (grant no. JP20bm0704046 to S.S. and T.S.) (grant no. JP18dm0207036 to H.B.), the Japan Society for the Promotion of Science (JSPS) (KAKENHI grant number 17H06312 to H.B.), and the General Insurance Association of Japan (the Medical Research Grant 2018 to K.A.)

## Author contributions

K.A., N.N., K.I., T.K., M.S., S.S., J.K., M.N., and H.O. designed the experiments. K.A., T.K., M.K., K.K., R.S., Y.K., S.S., and T.S performed the experiments. S.I., A.M., M.O., H.B. and K.K. provided the plasmids or viral vectors and contributed to interpreting the results. K.A. and all the other authors prepared the final manuscript.

## Competing Interests

M.N. declares a consultancy role with K-Pharma and research funding from RMic, Hisamitsu. H.O. declares a leadership position at the Keio University Graduate School of Medicine and is a compensated scientific consultant for San Bio Co., Ltd, and K Pharma Inc. All the other authors declare no competing financial interests.

---

**Figure 1–figure supplement 1.**
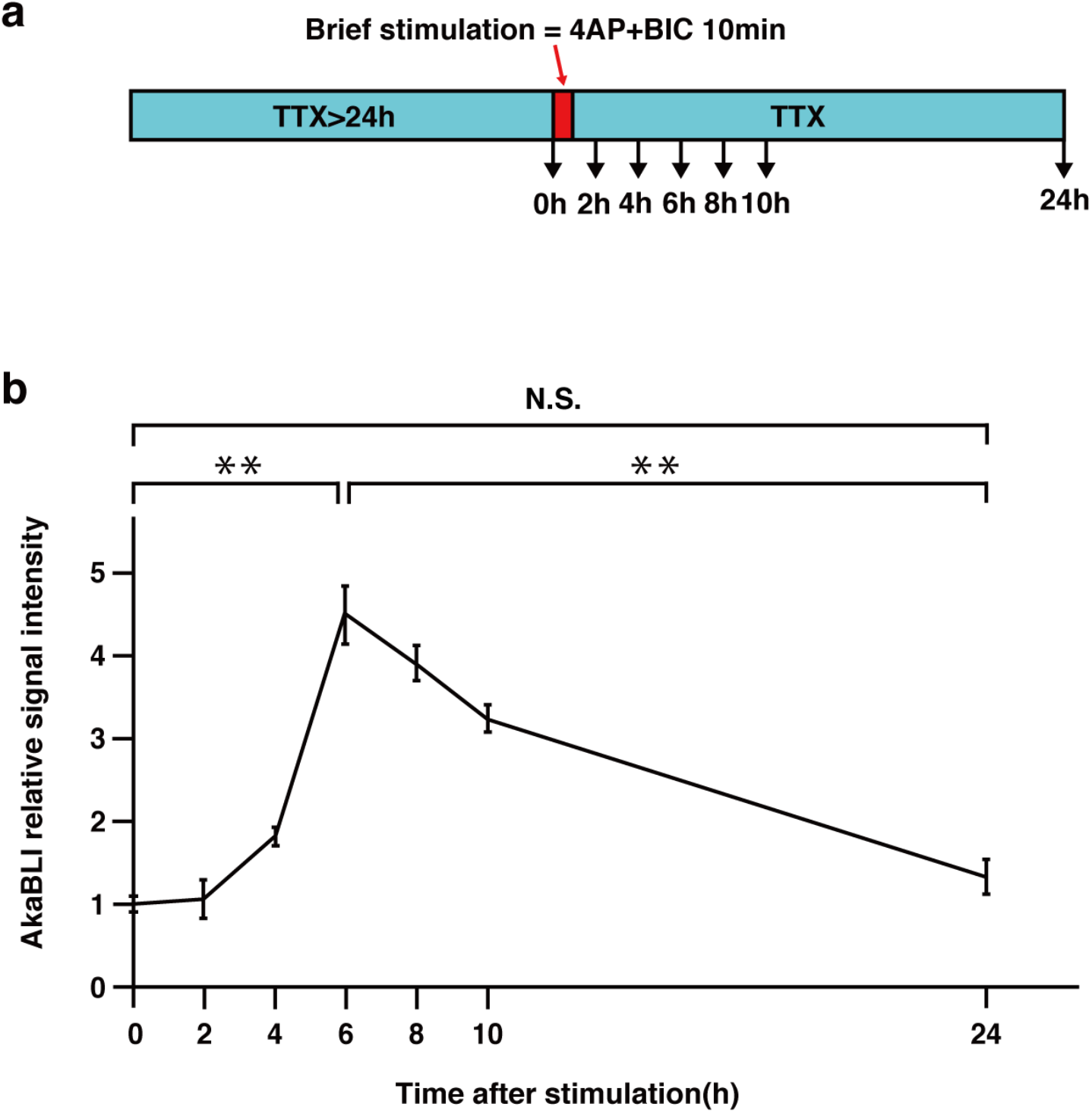
BLI of hippocampal neurons stimulated with 4AP+BIC *in vitro*. **(a)** Scheme of the experiment used to detect the time course of ESAL expression using mouse hippocampal neurons. Before and after brief stimulation, the neurons were silenced with 1uM TTX. **(b)** The result of multiple time-point BLI measurements in ESAL-expressing mouse hippocampal neurons. N = 5 independent experiments for 0, 6, 24 hours, and n = 3 for 2, 4, 8, 10 hours. Values are the mean ± SEM; *p < 0.05, **p < 0.01. Statistical analyses were performed using the two-sided unpaired Student’s *t* test. Individual t-values and degrees of freedom: 0 and 6 hours; t (8) =10.42, p= 6.2×10^-6^, 6 and 24 hours; t (8) = 8.357, p = 3.2×10^-5^, 0 and 24 hours; t (8) = 1.568, p = 0.16.

**Figure 3–figure supplement 1.**
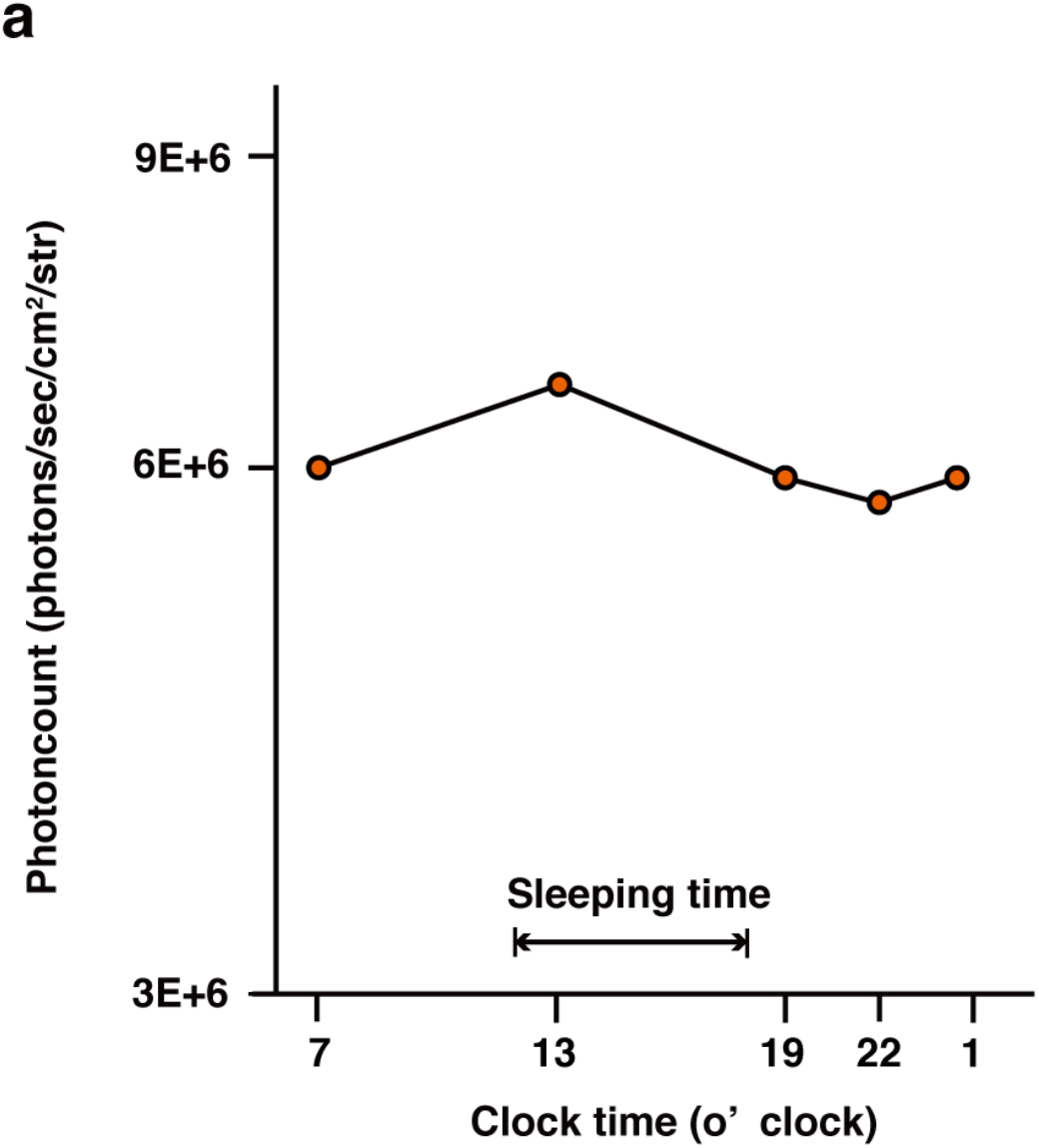
*In vivo* monitoring of ESAL bioluminescence over the day. **(a)** Representative diurnal transition of luminescence intensity in the cervical spine of an ESAL-expressing NS/PC-transplanted mouse (photons/sec/cm^2^/str). All the measurements were performed within a week (10–11 weeks after transplantation). Sleeping time in the figure shows a typical example because mice are nocturnal.

